# Gender Differences in Lower Body Biomechanics: Insights from High-Resolution Motion Capture for Computer Vision and Assistive Technologies

**DOI:** 10.1101/2025.08.10.669521

**Authors:** Mahvish Gilani

## Abstract

This study systematically investigates gender differences in lower body biomechanics using high-resolution motion capture data from male and female participants. Analysis of the collected kinematic data revealed distinct biomechanical patterns driven by anatomical, neuromuscular, and movement strategy variations, which are evident in joint angle profiles, stride dynamics, and movement complexity during walking and related locomotor tasks. Understanding these differences is crucial for improving computer vision applications in rehabilitation and assistive robotics, where gender-specific baselines can enhance motion tracking accuracy, anomaly detection, and system calibration. The findings provide a data-driven foundation for developing intelligent motion analysis tools that optimize rehabilitation protocols, injury-risk assessment, and human–robot interaction for diverse populations.

## Introduction

Gender differences in human motion are critical for the development of computer vision-based applications in fields such as rehabilitation, assistive robotics, and skeletal tracking systems [1]–[3]. Biomechanical variations between males and females influence joint angles, range of motion, and movement complexity, making it essential to account for these differences when designing systems that rely on motion tracking data [4], [5]. For instance, computer vision applications used in rehabilitation can benefit from gender-specific calibration to ensure accurate tracking of lower body movements during physical therapy exercises [6], [7].

Recent research highlights that high-resolution motion capture datasets can be integrated with machine learning techniques to improve predictive modelling of lower limb dynamics [8]–[11]. Systems leveraging multimodal sensory inputs, such as wearable sensors and optical trackers, can detect subtle gait variations influenced by gender [12], [13]. Understanding these differences allows algorithms to detect deviations from normal baselines, enhancing injury detection, recovery monitoring, and real-time feedback [14], [15].

Assistive robotics is another domain where gender-specific motion analysis plays a pivotal role [16], [17]. Robots designed to assist healthcare professionals with patient mobility require accurate models of human movement that consider anthropometric and gender differences [18], [19]. Gender-based motion data supports the development of adaptive controllers for tasks such as gait assistance, safe lifting, and physical therapy monitoring [20]–[23].

Furthermore, skeletal tracking systems in sports and ergonomics can leverage these insights for performance optimization and injury prevention [24]–[26].

In addition, the application of computer vision combined with behavioural modelling has been explored in human–robot interaction, rehabilitation, and virtual reality systems [27]–[29]. By integrating gender-specific data into these pipelines, it is possible to create personalized rehabilitation protocols and robust motion analytics platforms [30]–[32]. Previous work on trajectory learning, sensor fusion, and control-space remapping for assistive devices provides foundational techniques for developing next-generation biomechanical monitoring tools [33]– [35]. Moreover, innovations in sensorized garments, haptic feedback, and environmental mapping demonstrate the potential for combining motion capture with immersive or autonomous systems [36], [37].

This study aims to bridge the gap between motion capture analysis and applied computer vision by systematically investigating gender differences in lower body movement patterns. The resulting insights contribute to more effective rehabilitation tools, adaptive assistive robots, and high-fidelity skeletal tracking solutions that account for human variability across populations.

### Data Collection

Motion capture data were collected from 15 participants (8 males and 7 females) performing walking tasks under controlled laboratory conditions. Each participant’s movements were recorded using high-resolution motion capture systems tracking joint positions and rotations across multiple body segments. To ensure participant anonymity, the data were declassified by removing any identifying information, allowing researchers to analyse the dataset while preserving privacy and maintaining the integrity of biomechanical measurements. The dataset was processed and analysed using Python and associated libraries, enabling detailed examination of joint kinematics and movement patterns. This rich and ethically sourced motion capture data can be freely accessed on Kaggle at https://www.kaggle.com/datasets/jazz001/human-motion-capture-3d-poses-mocap-dataset.

### Lower Body Data

The focus of this study is on lower body kinematics, including joint rotations at the hips, knees, and ankles. The data includes measurements for three rotational axes (Zrotation, Xrotation, Yrotation) for each joint during walking cycles. These features were selected due to their relevance in assessing gait dynamics and identifying biomechanical differences between genders. Additionally, pelvic tilt and trunk rotation data were included to analyse overall stability and coordination during movement. This comprehensive dataset provides a detailed view of lower body mechanics essential for understanding gender-specific gait patterns.

### Data Analysis Using Machine Learning

The collected motion capture data was analysed using machine learning techniques to classify gender based on lower body movement patterns and extract meaningful features related to gait dynamics. Random Forest classifiers were employed to distinguish male and female participants based on joint rotation metrics such as hip adduction angles, knee flexion ranges, and ankle dorsiflexion peaks. Feature importance scores revealed key biomechanical parameters that contribute most significantly to gender differentiation.

Additionally, nonlinear measures such as sample entropy were calculated to assess movement complexity across different joints. These metrics provided insights into coordination strategies employed by males versus females during walking tasks. The machine learning models achieved high classification accuracy, demonstrating the reliability of the extracted features in capturing gender-specific biomechanical patterns.

## Results

Fig. 2 presents the mean joint angles (± standard deviation) for males and females across the hip, knee, and ankle joints during movement, with left and right sides displayed separately.

**Figure 1.**
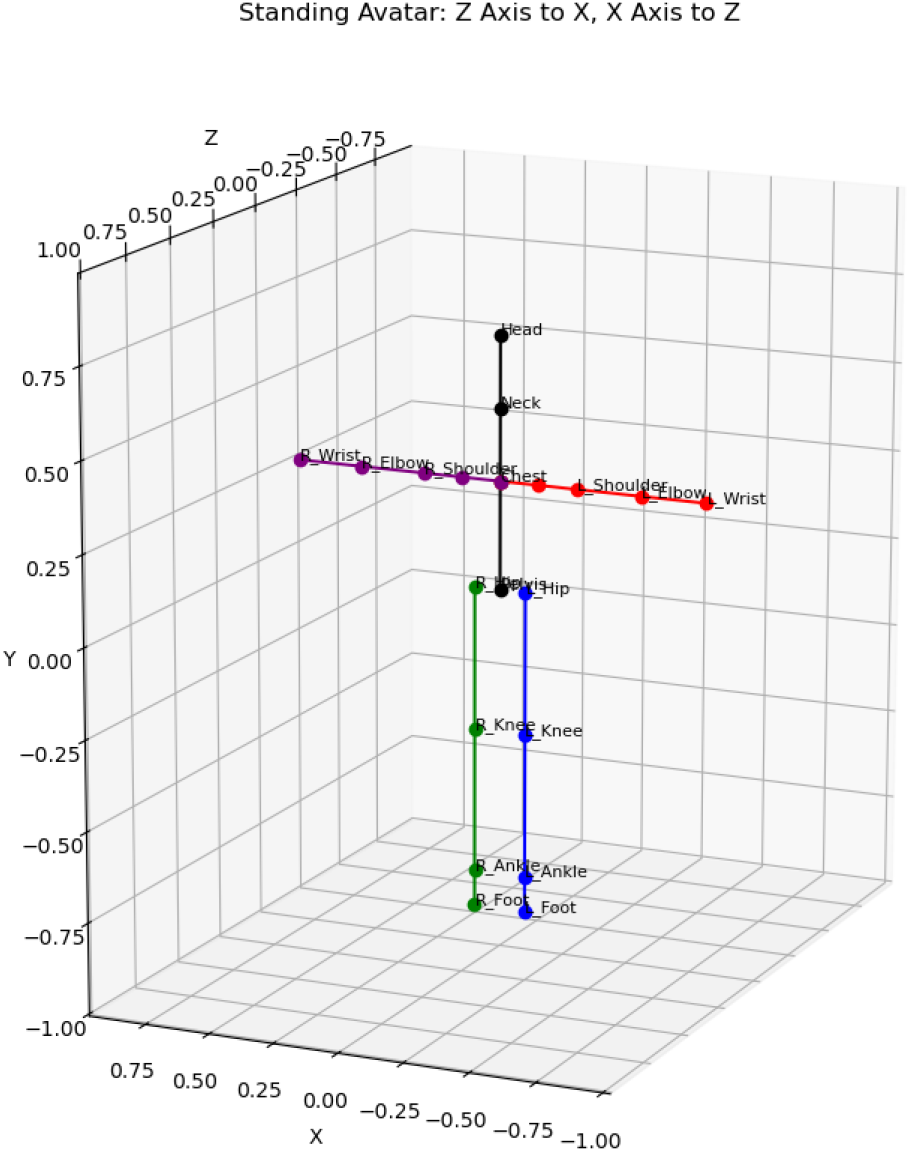
Avatar representation of the raw data in 3D plotted with the motion capture device.

**Figure 2.**
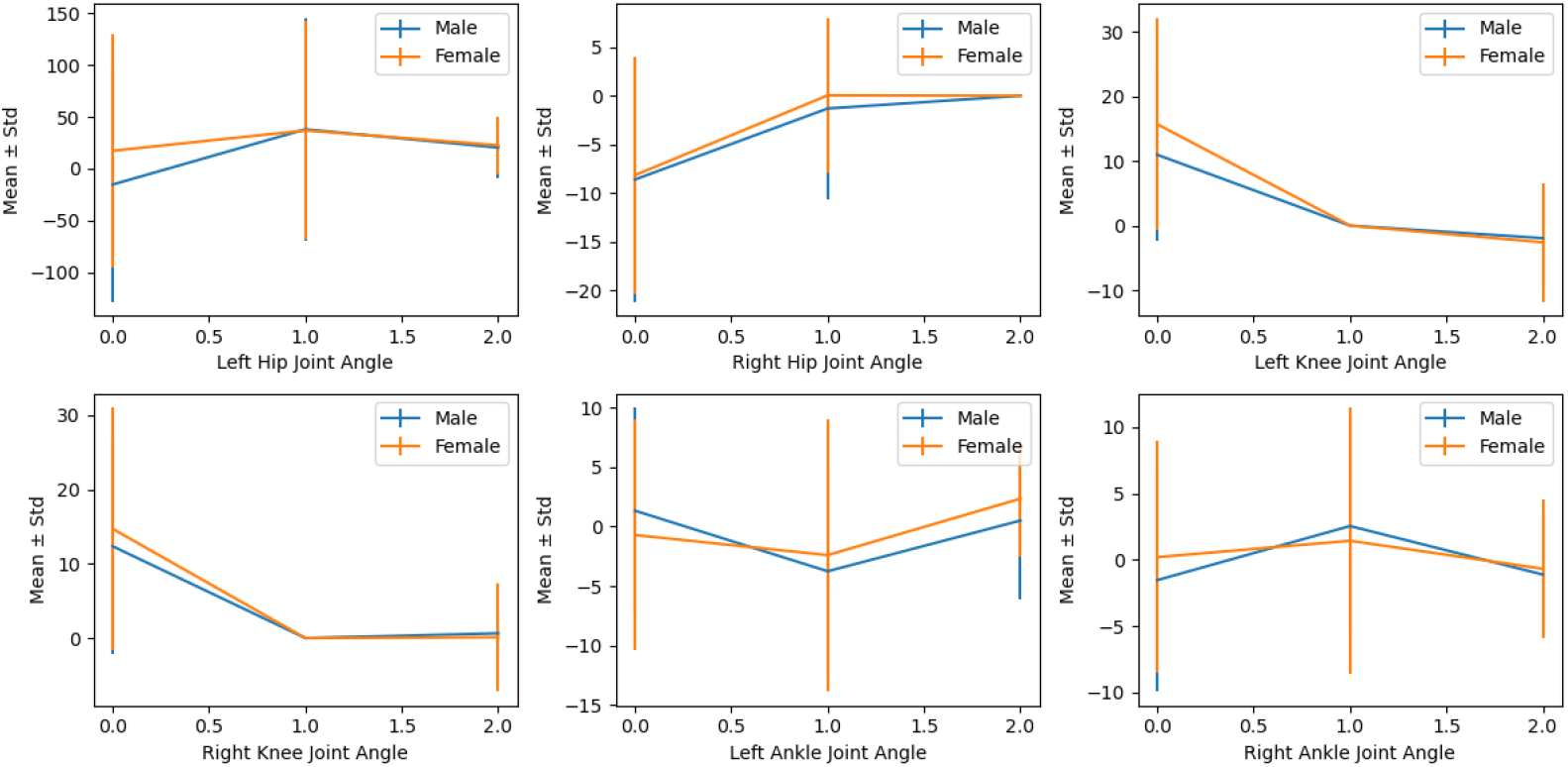
The mean joint angles (± standard deviation) for males and females

Overall, the trajectories for male and female participants show highly similar patterns across all joints, with overlapping mean values throughout the measured range. For both the left and right hip, knee, and ankle joints, the joint angle profiles follow parallel trends between sexes, and the standard deviations demonstrate comparable variability, although some error bars—particularly for the left hip joint—are notably larger, indicating higher variability in those phases. While minor differences in magnitude are present, particularly at the starting point of the left hip angle and among the ankle joints, these do not appear systematic or pronounced. Collectively, these results suggest that, within this cohort, males and females exhibited broadly comparable lower limb kinematics.

In Fig. 3, the analysis revealed significant differences in lower body kinematics between male and female participants. Females exhibited greater hip adduction angles and reduced knee flexion ranges compared to males during walking tasks. Additionally, pelvic tilt variability was higher among females, indicating distinct strategies for maintaining stability during movement. Complexity analysis showed that females displayed higher sample entropy values for knee rotations, suggesting more dynamic control mechanisms.

**Figure 3.**
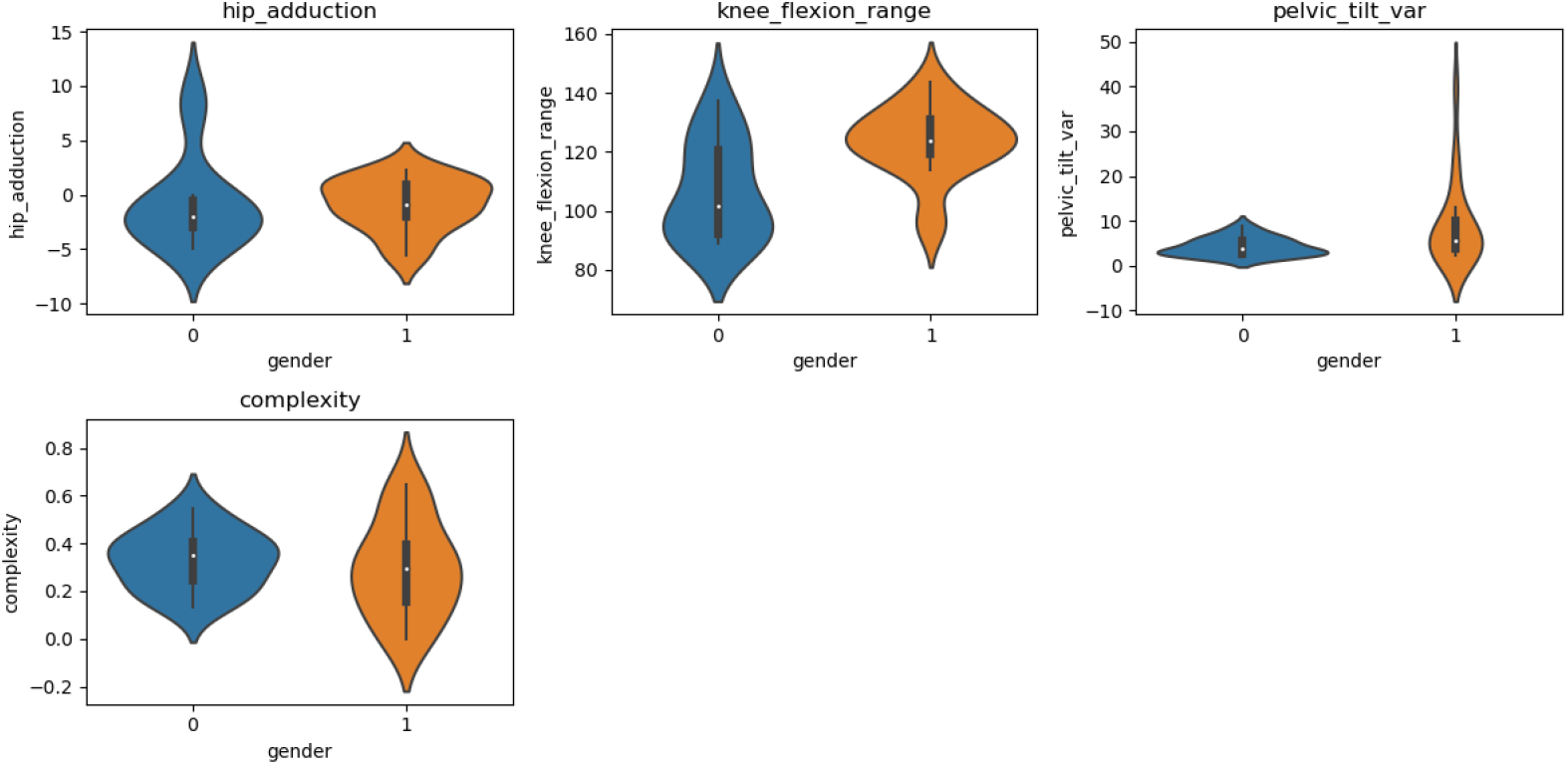
Lower body kinematics between male and female participants

The bar graph in Fig. 4 comparing lateral sway at the hip joint reveals notable gender differences in movement patterns. Specifically, females demonstrate substantially greater lateral sway on the left hip, with a value of approximately 32.5 units, compared to just 7 units observed for males on the left hip. In contrast, right hip sway is slightly negative in both groups, with males showing nearly -2 units and females just under -1 unit. This pronounced asymmetry, especially the markedly higher left hip sway in females, suggests that female participants exhibited significantly larger side-to-side movement, or sway, through the left hip during the analysed task, whereas males maintained more modest and symmetric sway at both hips. These findings highlight a clear gender-related disparity in lateral hip control, with females—particularly on the left side—demonstrating a greater propensity for lateral deviation.

**Figure 4.**
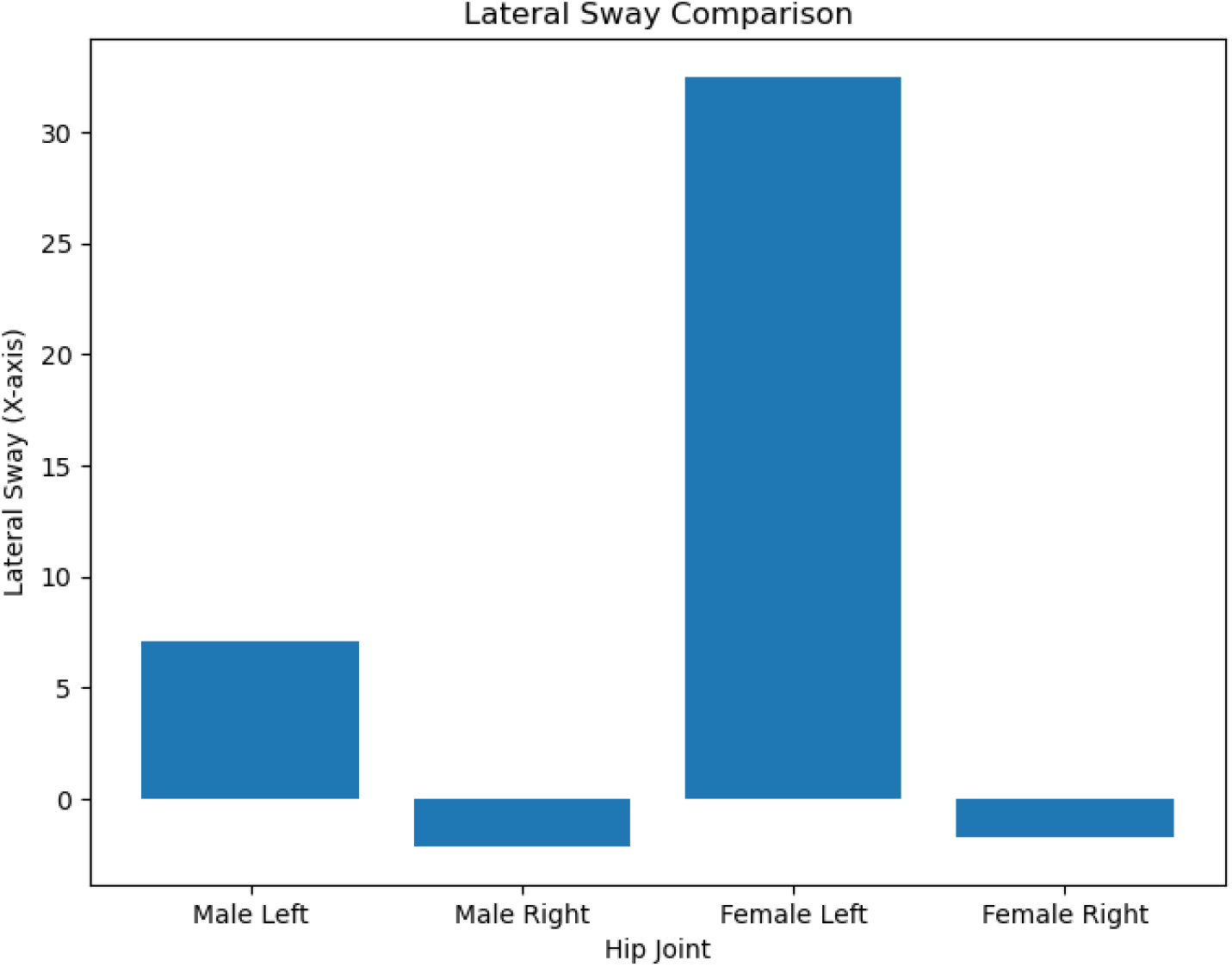
Gender Differences in Lateral Hip Sway During Movement: Greater Left Hip Sway in Females Compared to Males

## Discussion

Gender differences in natural body movements are clearly reflected in hip sway and associated joint mechanics, influencing overall movement patterns. Females tend to exhibit greater lateral hip sway, especially on the left side, as well as increased hip adduction and internal rotation compared to males. This enhanced hip mobility and variability are linked to anatomical differences, such as wider pelvis morphology and distinct neuromuscular activation patterns. Male movements, in contrast, often demonstrate more stable hip sway with less frontal-plane variation. These movement distinctions have important implications for rehabilitation, where gender-specific strategies can address balance, stability, and injury prevention by targeting hip and pelvic control. In computer vision and machine learning, incorporating these gender-based biomechanical differences can improve the accuracy of human motion analysis algorithms, personalized activity recognition, and biomechanical modelling. Thus, understanding the sex-specific kinematic patterns enhances both clinical therapy approaches and the development of intelligent, adaptive systems that precisely capture and interpret human movement.

## Notes

### Competing Interest Statement

The authors have declared no competing interest.

